# Engineering Controlled Microporosity in Liquid-Core Soft Compartments via Compositional Tuning of Aqueous Immiscible Systems for Facilitated Cell Migration

**DOI:** 10.1101/2025.06.11.659026

**Authors:** Raquel C. Gonçalves, Mariana B. Oliveira, João F. Mano

## Abstract

The processing of materials at aqueous interfaces has enabled the generation of compartmentalization structures with broad biomedical interest. Controlling the porous structure of soft materials to create micro-sized pores that enable cell migration is crucial for tissue development and regeneration. This work presents liquid-core soft compartments formed via interfacial polyelectrolyte complexation between alginate and ε-poly-L-lysine, which membrane physical properties are tailored by adjusting the composition of the prototypical aqueous two-phase system. The additional interfacial dynamics provided by the presence of the immiscible polymer phases promoted the organization of porous architectures in the membrane. Fiber-shaped tubular structures enable adhesion and migration of mesenchymal stem cells to surrounding fibrin matrices and its invasion, behavior that was significantly improved in conditions of higher membrane porosity. While preserving the single-step approach of the established technology, the interfacial materials with tailorable porosity can be processed for applications in tissue engineering and regeneration.

## 1. Introduction

The porosity of soft materials is crucial not only for ensuring adequate permeability and nutrient diffusion but also to promote the migration and invasion of cells.[1] Pores at the micrometer scale (1-100 µm) are required for the free movement of cells and dynamic cell processes, including tissue ingrowth, angiogenesis and cell transmigration.[1–4] Although some methods to achieve this scale of porosity have been developed, they are associated with some limitations. Most conventional approaches rely on sacrificial templates, or porogens such as polymeric microparticles, that need subsequential removal to create void spaces in the solidified material.[1,5] Also, freeze-drying[6] and cryogelation[7] methods are associated with either a reduction of the material’s density and consequent mechanical integrity, or limited control over pore uniformity. Alternatively, techniques such as bioprinting[8], photolitography[9,10] and laser ablation[11–13] offer the possibility to generate pores, holes or channels with better resolution; however, these methods are often time-consuming and/or require specialized equipment as well as qualified people. Liquid-liquid phase separation in polymeric systems has become a simpler strategy to generate porous hydrogels with diverse morphologies showing improved cell stretching and migration capacity.[14–17] While these techniques are predominantly used in bulk soft materials produced in continuous environments, the potential to tailor porosity specifically at interfaces has received limited attention.

In recent decades, the organization of macromolecules at interfaces has enabling developing chemically tailored surfaces and confining materials with wide interest. Assemblies and physical interactions can be promoted at solid-liquid interfaces, for example, by the sequential deposition of oppositely charged polyelectrolytes (PEs) on solid templates[18,19], or at liquid-liquid interfaces of miscible[20–22] or immiscible phases[23,24]. In fact, when two solutions of oppositely charged PEs are placed side-by-side, a soft membrane derived from the electrostatic interaction and complexation between the PEs is formed following the solution interface.[20] Additionally, when two immiscible liquids come into contact, interfacial tension arises to minimize the system’s free energy and maintain phase separation. If antagonistic charged PEs are introduced into the immiscible phases separately, they will spontaneously interact at the interface of two liquids.[25] Thus, in this case, two distinct interfaces and phase separation processes can be identified: one arising from the interaction between the PEs, and another resulting from the immiscibility of the two phases. Introducing a two-phase interface into the process may provide different driving forces for molecular interaction and assembly, enabling the possibility of manipulating the structure and architecture of the fabricated interfacial material.[26] Typically oil/water interfaces are used, but due to concerns over potential toxicity and the challenges of washing away oils or organic solvents, aqueous two-phase systems (ATPS) have gained attention as a biocompatible alternative for biomaterial production.[25] Soft membranes formed through PE complexation at all-aqueous interfaces, taking the form of spherical capsules or flexible tubular fibers, exhibit diverse features and functionalities interesting for a wide range of applications in the biomedical field, including the encapsulation and compartmentalization of biological cargo[27,28], the development of delivery vehicles[29], and the creation of reaction platforms[22,30]. In such cases, membrane permeability plays a crucial role in the regulation of molecules, materials or biological substances that can cross the membrane, which in turn is influenced not only by factors such as pressure differences, and the dimensions and surface characteristics of the membrane, but also its porosity.[1]

Here, we report a porogen-free and single-step method to increase the porosity of interfacial PE membranes, by adjusting the composition of the most classical prototypical polyethylene glycol (PEG)/dextran ATPS used to support the formation of tubular liquid-core fibers. In cell-laden membranous materials produced through all-aqueous interfacial complexation, cells are usually kept inside the designated compartment to promote self-assembly into microtissue aggregates[28,31–33], or to encapsulate single cells[34]. Alternatively, the membrane can act as a scaffold for cell adhesion, by incorporating cell-adhesive moieties or imparting a filamentous structure.[35,36] While these approaches show promise for organoid engineering or the development of tissue biomimetic models, the potential for highly porous aqueous compartments to promote the invasion of encapsulated cells into surrounding matrices remains unexplored. This could be relevant for biomedicine and tissue engineering fields given the critical role of cell migration and invasion in various biological processes including angiogenesis, wound healing and tissue regeneration.[37]

In this study, the interaction and complexation process between natural PEs, namely ε-poly-L-lysine and alginate, is observed to be influenced by the composition of the aqueous immiscible phases, thereby resulting in the spontaneous formation of distinct micro-structural porous features at the interfacial membrane. Interestingly, the overall macrostructure and mechanical integrity of the fiber structures remains unaffected by the enhanced microporosity of the membrane; and in fact, it improves their structural robustness. These properties are preserved independently of construct’s size and geometry, and demonstrate significant effects in the capacity to promote the migration of encapsulated mesenchymal stem cells and invasion of surrounding fibrin hydrogels. The developed method offers a simple and effective approach to modulate the morphological features of all-aqueous interfacial membranes, with potential applications in tissue engineering and regeneration.

## 2. Results

### 2.1. Fabrication of tubular fibers with different membrane porosity

Aqueous solutions of PEG (MW 8 kDa) and dextran (DEX, MW 450–650 kDa) were used as a prototypical ATPS to create a water/water interface, that control the interaction of oppositely charged PEs, each distributed separately in one of the phases, as previously reported.[28,35,38] Alginate (ALG, MW 100-200 kDa) and ε-poly-L-lysine (EPL, MW 4.7 kDa) in the dispensing and bath phase, respectively, diffuse and interact electrostatically at the interface of the solutions to form a coacervate-like membrane. The formation of easy to handle and macroscopically surface-homogeneous tubular fiber-shaped materials was achieved by depositing a DEX + ALG solution in a PEG + EPL bath phase, and applying continuous displacement of either the dispensing or matrix setups (Schematic representation in **Figure 1**A). Appropriate concentrations of both phase-forming or charged polymers were used considering previous optimized formulations that enabled the formation of physiologically stable and cytocompatible structures, namely, PEG 17 wt%, DEX 15 wt%, ALG 2 wt% and EPL 0.75 wt%.[35] After 2 minutes of complexation, the interface of the ATPS precursor is disrupted by performing minor washing steps, and structures are retrieved to a physiological saline medium.

**Figure 1.**
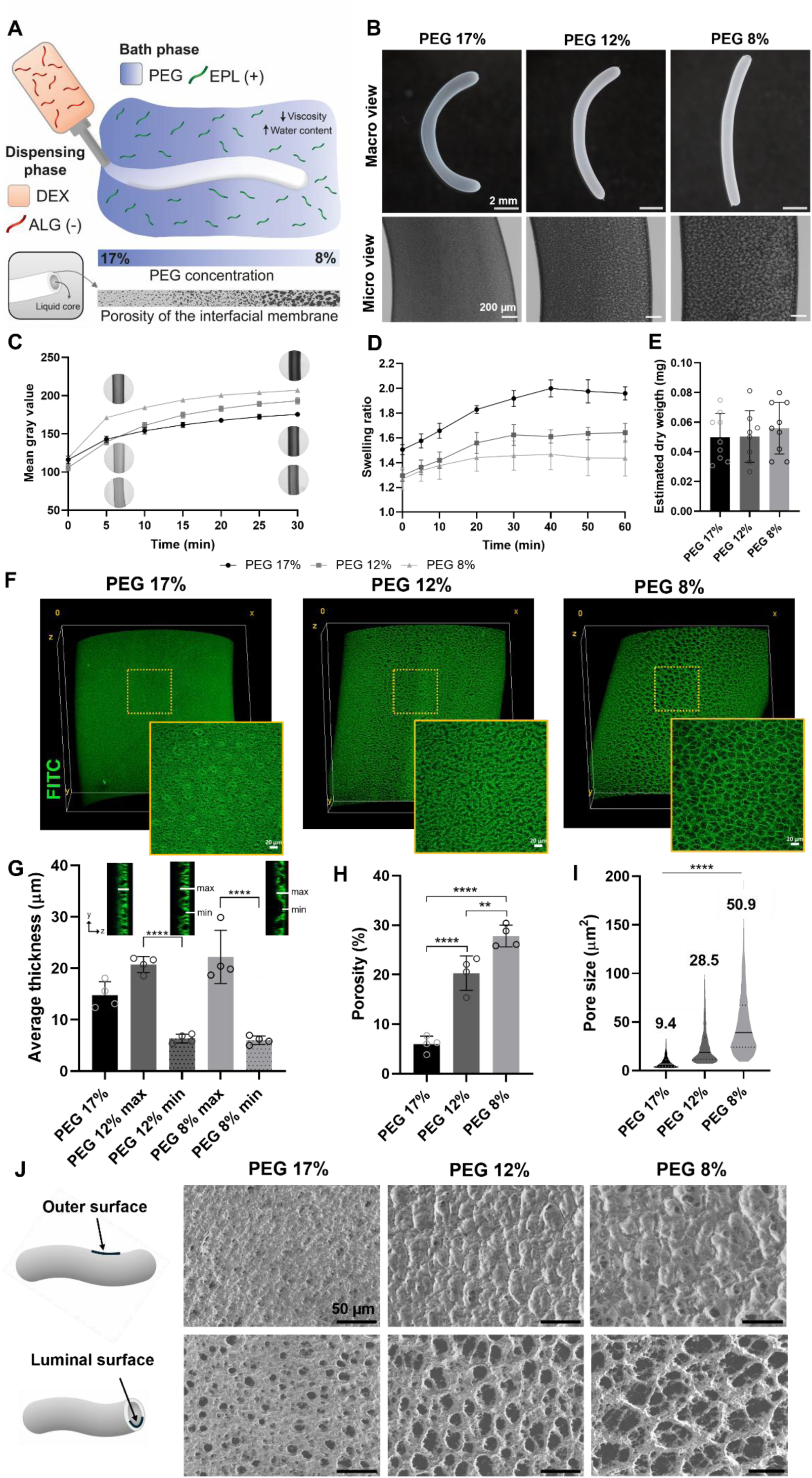
Fabrication of interfacial fiber-shaped membranes with diverse porosity and its structural characterization. **A** Schematic showing the fabrication of liquid-core fibers by dispensing a solution of dextran (DEX) and alginate (ALG) in a bath containing polyethylene glycol (PEG) and ε-Poly-L-lysine (EPL). Driven by electrostatic complexation between the ALG and EPL, an interfacial membrane is created. The porosity of the interfacial membrane is increased by adjusting the composition of the bath phase, namely, by decreasing the concentration of PEG, which provides a less viscous environment with a higher water content. **B** Representative macroscopic images of fibers in PBS produced in baths with different PEG concentration – 17%, 12% and 8% - as well as brightfield microscopic images. **C** Mean gray value over time of fibers produced in the different baths, indicative of fiber opacity (n = 4 replicate fibers). **D** Swelling ratio determined by external diameter variation over time after washing with PBS (n = 4 replica fibers). **E** Estimated dry weight after lyophilization (n = 8 (12%) or 9 (17%, 8%) replica plastic tubes containing fibers of the same length). **F** Three-dimensional representations of confocal micrographs of the interfacial membranes with hollow liquid cores, as well as closed images to show their porous morphology. Fibers were incubated with fluorescein isothiocyanate (FITC) for membrane visualization. **G** Average thickness measured from the orthogonal confocal projections. Maximum and minimum values were determined for the more heterogeneous membranes, namely the ones with lower PEG concentration – 12% and 8% (n = 4 replica fibers, 4 measurements per fiber). **H** Membrane porosity determined from z-stack maximum projections (n = 4 replica fibers). **I** Pore size distribution, with mean values represented at the top of each plotted condition (pores from n = 4 replica fibers). **J** Schematics and representative SEM micrographs of the outer and luminal surfaces of the fiber membranes. The structures were dehydrated with ethanol solutions with increasing concentrations. Data is presented as mean ± standard deviation (s.d.). Statistical analysis was performed using one-way ANOVA statistical analysis with multiple comparisons. **p<0.01 and ****p<0.0001.

In here, we carefully studied the possibility of tailoring the porosity of the interfacial membrane within the same system, by adjusting the concentrations of PEG and DEX (Figure S1, Supporting Information). While decreasing PEG concentration from 17 wt% to 2-8 wt% induced the formation of microscopic porous-like morphological features in the membrane, adjusting the DEX concentration did not produce such visible effects. Besides concentration, polymer molecular weight also strongly influences the density and viscosity of the solutions.[25] Due to the significant differences in molecular weight between the polymers (PEG vs. DEX), and the viscosity-driven effect of dissolved alginate, the dispensing solution exhibits higher density and viscosity compared to the bath phase. Therefore, we evaluated the effect of altering the properties of the prototypical dispensing phase, by using DEX with different molecular weights (MWs) (Figure S2, Supporting Information). Reducing the MW to 75 and 6 kDa resulted in interfacial fiber membranes with very distinct morphological characteristics. When using polymers with MW of the same order of magnitude, namely PEG 8 kDa and DEX 6 kDa, the membrane is poorly porous both before and after washing, while the use of DEX 75 kDa produced membranes with branched complexes with an organized interspacing after washing. Moreover, in the tested conditions, the reduction of the PEG concentration continued to induce the generation of more interspaced complex networks in the interfacial membrane, although after the washing steps this was more evident with the original 450-650 kDa DEX. This suggests that a large difference between the MW of the phase-forming polymers is necessary to cause a significant effect on the membrane porosity of the studied system. Although manipulating the properties of the prototypical ATPS shows potential for controlling membrane characteristics, it is important to consider its impact on the processing and handling of the resulting tubular fibers (Figure S3, Supporting Information). Very low polymer concentrations primarily hindered fiber processing due to decreased bath viscosity or density of the dispensing phase, while lower DEX MW led to fibers with reduced handling quality.

To determine whether a specific ATPS polymer plays a key role in forming interfacial tubular membranes with enhanced porosity, we empirically investigated the impact of removing each polymer from the process. Similar morphologies were visualized after the complete removal of PEG from the bath phase, although the membrane is less openly porous (Figure S4, Supporting Information). Moreover, tubular membranes formed with a dispensing phase containing only ALG (without DEX), exhibited a more homogeneous and less dense structure. Both these observations highlight an essential role of the DEX phase in generating the pore-like microstructural changes in the membrane (Figure S4, Supporting Information). However, only when both the polymer phases are present, the global porosity of the membrane is significantly affected. In membranes formed in conditions where both of the polymers are absent, very dense interfacial coacervates with no structural organization are formed (Figure S4, Supporting Information).

Since the concentration of PEG played a crucial role in achieving desirable membrane porous features, we selected three formulations in which only the PEG concentration varied, namely 17 wt% (used as control), 12 wt% and 8 wt%, to conduct a more detailed study and characterize membrane porosity. Macroscopically all fiber structures are similar with no evidence of increased porosity in the membrane, but microscopic observations revealed pore-like heterogeneities in the membrane as the concentration of PEG decreases (Figure 1B). The gradual increase in opacity (measured by the mean gray value) with time correlates with the timely diffusion and continuous interaction of the PEs, thereby generating fiber structures with increased membrane thickness and robustness.[35] Capito and colleagues demonstrated that the complexation between large and small oppositely charged molecules resulted in interfacial membranes that grow in a highly ordered architecture, given the osmotic pressure imbalance that influenced the diffusion of the PEs and their self-assembly process.[21] In here, the process of membrane growth is believed to be similar considering the different affinity of the PEs to the ATPS polymer phases. Previous studies on PEs partition coefficient revealed that while ALG strongly prefers the DEX phase, EPL distributes between both aqueous phases, with a higher affinity for the inner DEX phase, opposite from where it is originally dissolved in the process of fiber formation.[35] This suggested that EPL diffusion is the primary factor driving membrane formation, and resulted in an inward growth of the membrane at extended complexation times. The higher degree of opacity in fibers produced using PEG at 8% right after 5 minutes of complexation, indicates an accelerated and favored PE interaction (Figure 1C). After 30 minutes, differences in the gray value demonstrate the effect of decreasing the PEG concentration, leading to progressively darker membranes. Since less viscous environments facilitate molecular diffusion, a greater number of polyions are expected to interact due to the faster and freer movement of EPL chains towards the interface at reduced polymer concentrations.

Upon washing with PBS, a clear swelling of the fibers was observed due to osmotic effects caused by the disruption of the prototypical interface.[35] This swelling was evaluated as a function of time, considering the external diameter variation during volumetric fiber expansion. While in PEG 17% formulation, a gradual increase in swelling ratio is observed over time, 12% fibers exhibited less expansion, with minimal swelling in fibers produced with 8% (Figure 1D). Moreover, the time required to reach equilibrium decreased from 40 to 20 minutes due to the lower PEG concentration. This demonstrates an enhanced mechanical robustness of the membrane in PEG 8% formulation, allowing it to better withstand water-swelling effects and maintain its structural dimensions over time. This behavior could be explained by the accelerated complexation, and an apparent densification of the interfacial membrane. Nevertheless, interestingly, the dry weight of fibers of 0.5 cm length was estimated at around 50 µg consistently across all three conditions, with no significant statistical differences observed (Figure 1E).

To characterize the morphological properties of the membranes, we labeled the fibers with fluorescein isothiocyanate (FITC) and examined them across the membrane thickness using confocal fluorescence microscopy (Figure 1F). The thickness, determined from orthogonal projections, slightly increased from ∼14 µm to a maximum of ∼22 µm (Fig. 1H). However, given the higher porous nature of the 12% and 8% membranes, we determined a minimum thickness of approximately 6 µm in the pore regions. The membrane porosity significantly increased from approximately 6% to 28% as the PEG concentration decreases from 17% to 8%, respectively (Figure 1H). Regarding the size of the pores, we mostly consider their area given their irregular shape (Figure 1I). Membranes produced in baths of variable PEG concentration have a mean pore size of 9.4, 28.5 and 50.9 µm^2^, with average Feret’s diameter of 5.1, 9.4 and 11.8 µm, for the respective 17%, 12% and 8%. Scanning electron microscopy (SEM) was used to evaluate the surface characteristics of the outer (facing the external environment) and inner (facing the lumen of the tubular fibers) parts of the membranes. SEM micrographs indicate that the porous morphology is mostly present in the internal surface of the membranes, while the outer surface is more homogeneous, and becomes rougher when PEG concentration is decreasing (Figure 1J). Moreover, closer images of the pores at PEG 8% indicated an interesting highly filamentous organization (Figure S5A, Supporting Information). Together, these observations suggest that the pores in the lower concentration formulation are covered by a very thin outer layer that is more amorphous, which is the result from the initial interfacial PE interaction at the early stages of liquid-liquid contact. As the smaller EPL chains (compared to ALG) diffuses towards the inner liquid, the membrane grows and forms an inner thicker filamentous layer that is organized with pore-like heterogeneities, as the result of the electrostatic self-assembly and a possible structural rearrangement in the presence of the immiscible components. Moreover, time-lapse videos of the z-stacks of PEG 8% membranes, reveal a noticeable increase in pore size throughout the membrane thickness from the outer region to the inner lumen, indicating an additional heterogeneity and hierarchy through the z axis of the membrane structure (Video S1 and Figure S5B,C, Supporting Information). This asymmetry aligns with the observations of Baig *et al.* [39], and can be attributed to the dynamic complexation process in which the rate of EPL entry varies during membrane formation.[38]

To explore the versatility of the purposed method, we investigated the effect of other parameters, such as complexation time and the size and geometry of the final compartment, on the porosity of the membrane, while maintaining PEG at 8%. Higher complexation times produced membranes with lower pore sizes, with mean values of 40.3 µm^2^ and 26 µm^2^ for 15 and 30 minutes of complexation, respectively (Supplementary Fig. 6). On the other hand, membrane porosity is maintained, showing a slight increase, across liquid-core fibers with diameters ranging from 0.47 to 2 mm, produced with needles of 27 to 20 gauge, respectively (**Figure 2**A,C). Additionally, we observed very similar porous structures in spherical-shaped membranes, of both small and large sizes, with diameters of 0.8 mm and 5 mm, respectively (Figure 2B,C). In general, the results show a tendency for decreasing porosity and pore size with the radial dimension of structures, whether they are fibers or spheres. The pore size decreased to mean values of approximately 40 µm^2^ for the smallest structures compared to values within the range of 90 to 100 µm^2^ for the larger ones (Figure 2D).

**Figure 2.**
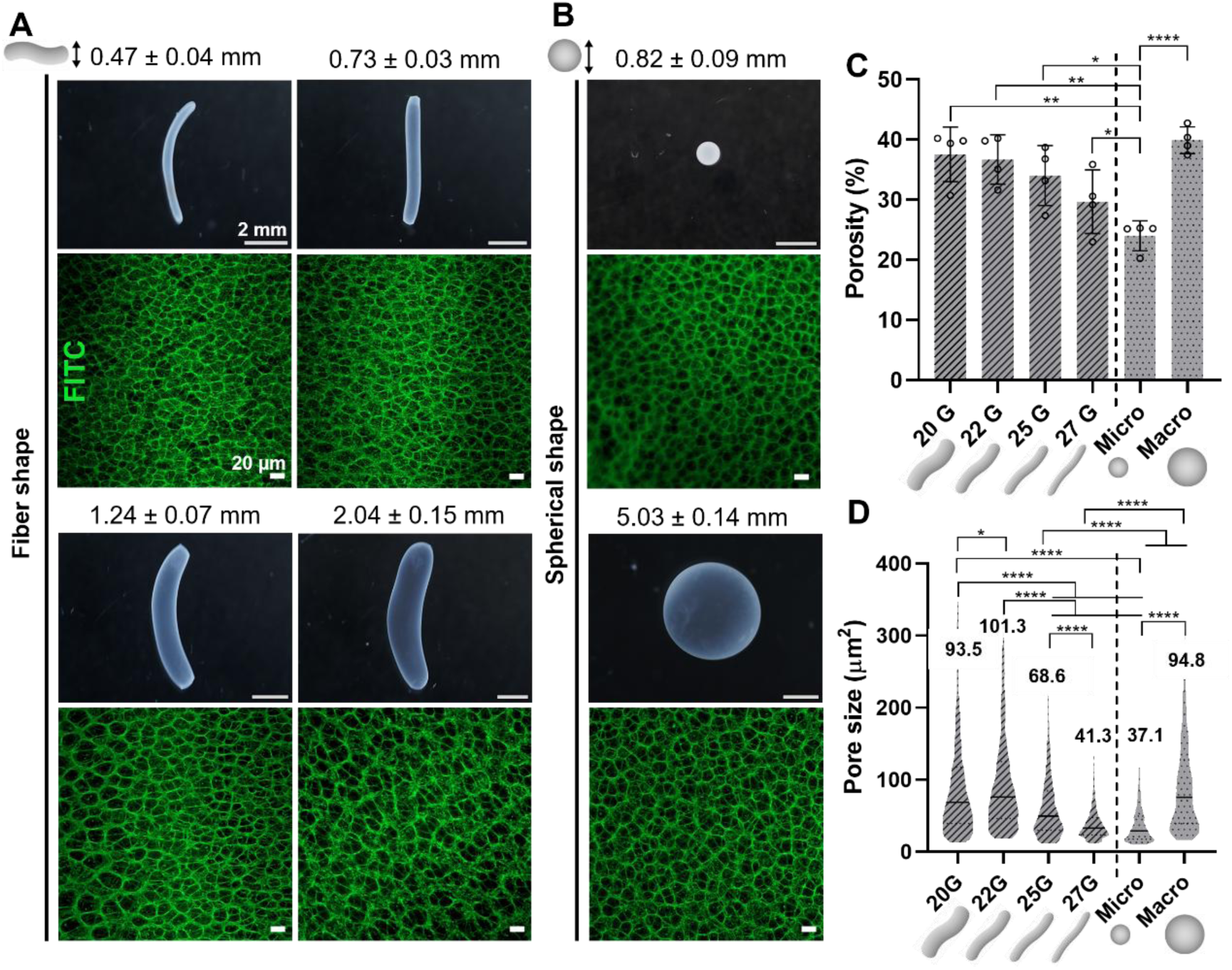
Influence of the size and shape of the final compartment on membrane porosity. Photographs (top) and confocal fluorescent micrographs (bottom) of washed fiber-shaped in **A** and spherical-shaped in **B** compartments with variable sizes. Mean and standard deviation values of their external diameter are presented for each fiber and sphere size. **C** Quantification of membrane porosity from the confocal maximum projection images (n = 4 replicas for each condition). Data are mean ± s.d. with *p<0.05, **p<0.01 and ****p<0.0001; one-way ANOVA with multiple comparisons. **D** Pore size distribution with mean values represented at the top of each plot (pores from n = 4 replica fibers).

### 2.2. Cell-mediated liquid-core fiber deformation

The potential effect of the membrane’s porous morphology on cell behavior was assessed by directly mixing human adipose-derived mesenchymal stem cells (hASCs) in the dispensing phase, that also contained ALG functionalized with the arginine-glycine-aspartic acid sequence (ALG-RGD) to promote cell adhesion to the membrane. This phase was dispensed in bath solutions containing PEG 17% and 8% as the control and more porous conditions, respectively. After producing the cell-laden fibers, they were washed, transferred to cell culture medium and cultured for 7 days, for subsequent viability analysis. Live/dead assay revealed that reducing PEG concentration didn’t affect cell viability, with cells remaining highly viable in both conditions along the culture period, indicating the cytocompatibility of the method (**Figure 3**A). At day 1 there is an apparent less aggregation of the cells in the PEG 8% membranes. This tendency was also visualized in mouse endothelial cell line, but throughout the 7-day culture period (Figure S7, Supporting Information). GFP-expressing C166 cells were more uniform distributed within the fibers of PEG 8% formulation, while they formed larger aggregates in the control condition. This suggests that the characteristics of the porous structure of the interfacial membrane may distinctively affect the distribution patterns of different cell types within the compartment. The hASCs started to adhere right after 1 day and maintained a spread morphology along the 7 days, while they continuously proliferate inside the fiber structure with no statistical differences on cell metabolic activity between conditions (Figure 3B).

**Figure 3.**
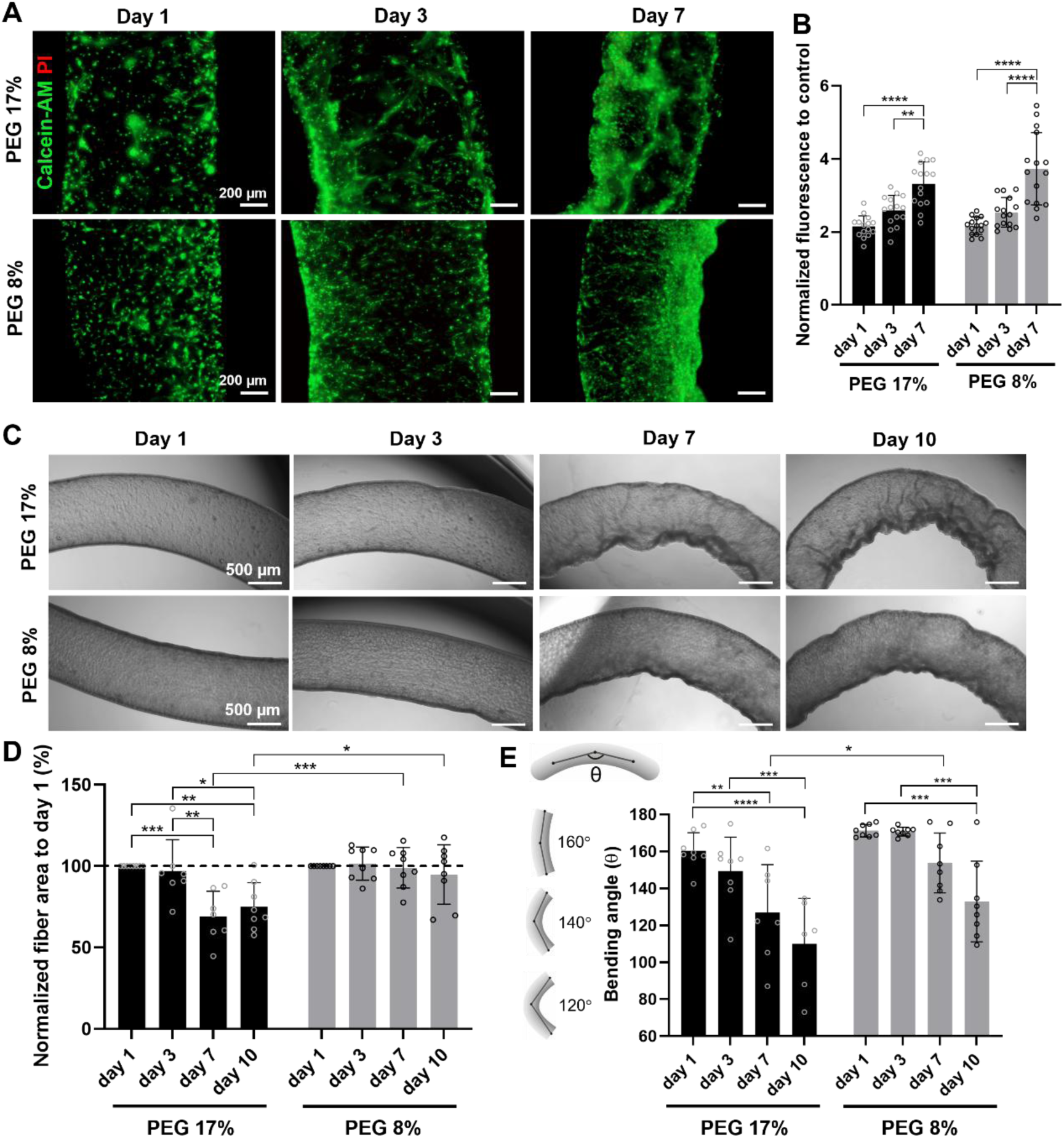
Cell-laden fibers and their ability to undergo deformation. **A** Live (calcein-AM)/dead (propidium iodide, PI) micrographs of human adipose-derived mesenchymal stem cells (hASCs) encapsulated within the fibers after 1, 3 and 7 days of culture. **B** Cell metabolic activity plotted as the fluorescence intensity of cell-laden fibers normalized to control fibers without cells, using AlamarBlue^TM^ assay (n = 15 replica fibers from 3 biologically independent experiments). **C** Representative optical microscope images of cell-laden fibers, throughout 10 days of culture. **D** Quantification of fiber area reduction over time compared to day 1 (n = 8 replica fibers from 2 biologically independent experiments) **E** Quantification of the bending angle (θ), measured at the center of each fiber as is represented by the schematic, from microscope images at the different timepoints (n = 8 replica fibers from 2 biologically independent experiment; eventual fibers that didn’t undergo the prevailing bending were excluded). Mean ± s.d., two-way ANOVA statistical analysis with multiple comparisons. *p<0.05, **p<0.01, ***p<0.001 and ****p<0.0001.

During cell culture, we observed that cells were folding the membrane, causing a macroscopic deformation of the fiber structures. The ability of human MSCs to fold materials was previously observed in soft fibrous hydrogels and thin films, leading to the formation of dynamic constructs with spontaneous deformation or the development of spheroidal structures with minimized necrotic core formation.[40–42] Fiber deformation was assessed by the reduction of the fiber area throughout 10 days of culture, determined from optical microscope images (Figure 3C). A significant decrease in fiber area occurred in the PEG 17% condition after 7 and 10 days, with fibers contracting down to ∼75% of their original area at day 1 (Figure 3D). This behavior was not observed in the morphologically more porous fibers produced with the PEG 8% bath (Figure 3D). In addition, a tendency for them to undergo bending into more favorable geometries was observed in most cases, mainly due to gravity-driven effects that promoted an increased cell density at specific regions of the fiber compartment, gradually forming more curved structures over time. By measuring the bending angle, which corresponds to the vertex angle at the center of each fiber[43], there was evidence of a stronger bending deformation in PEG 17% fibers after 7 days, compared to the PEG 8% (Figure 3E). These findings support the increased robustness of membranes in the PEG 8% formulation, making them more resistant to deformation. The contractile forces of hASCs were believed to be the main driving factor for this behavior. Cell contractility is mediated by the aggregation of the motor protein myosin II, and its interaction with overlapping actin filaments, leading to cytoskeletal tension.[44] As such, we evaluated the effects of suppressing contraction using an inhibitor of myosin II – blebbistatin – which was previously shown to promote a distinct cytoskeleton organization with thinner filaments in hASCs, hampering their adhesion capacity.[45] The treatment of cell-laden fibers with blebbistatin at day 3 prevented membrane folding and fiber deformation, with no reduction on fiber area after 7 days of culture, in both conditions (Figure S8A,B, Supporting Information). Additionally, treating 7-day deformed fibers resulted in slight de-folding behavior after 1 day with a small increase in the bending angle although not statistically significant, and further maintenance of fiber geometry over 7 days, in both conditions (Figure S8A,C, Supporting Information). Altogether, these results indicate the distinct ability of the soft compartment to enable adhered cells to plastically deform the membrane through contraction resulting in interesting fiber geometries, that are totally dependent on the cell’s behavior and membrane properties.

### 2.3. Cell migration from fiber compartments to surrounding matrices

Cell migration in a material depends on cell-cell and cell-matrix interactions, where specific biochemical and biophysical cues, such as growth factor signaling and material porosity, respectively, play a major role in stimulating/guiding cell motility across the material. Several studies have been conducted to assess the material’s ability to support cell migration, primarily by embedding spheroids and analyzing cell outgrowth[46], ones reporting the effectiveness of highly porous hydrogels to potentiate cell migration.[47] In here, we hypothesize that the higher porosity and pore size of the PEG 8% membranes may enhance the ability of encapsulated and adhered cells to migrate from the fiber membrane to a surrounding matrix, compared to the less porous and uniformly thick PEG 17% membranes. To explore this hypothesis, cell-laden fibers in which cells were left to adhere to the different membranes for 4 days, were embedded within fibrin hydrogels as schematically represented in **Figure 4**A. Fibrin is a natural protein formed during blood clotting, through the enzymatic cleavage of fibrinogen by thrombin, leading to the polymerization of insoluble fibrin monomers into a three-dimensional network.[48,49] Fibrin hydrogels have been widely used in cell migration assays due to their biomimetic properties, closely resembling the extracellular matrix.[50] They support cell adhesion, remodeling, and proteolytic degradation, effectively replicating physiological migration in wound healing and tissue regeneration.[48,50]

**Figure 4.**
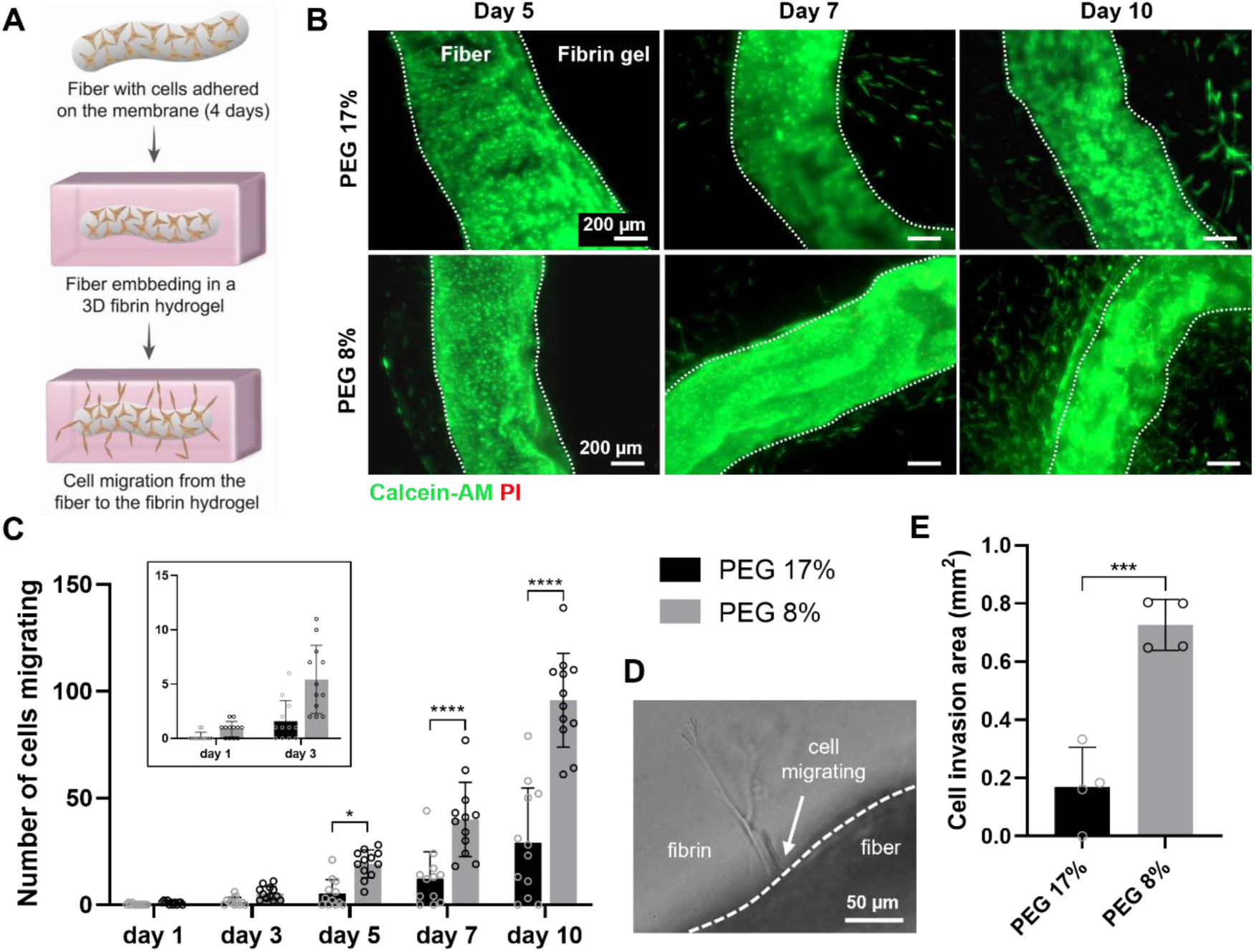
More porous interfacial membranes facilitate the migration of adhered cells and invasion of surrounding hydrogels. **A** Schematic representation of the method used to study the migration of the cells, by embedding cell-laden fibers with 4 days of culture to promote cell adhesion to the membrane, within fibrin hydrogels. **B** Representative live (calcein-AM)/dead (PI) fluorescence microscope images of cell-laden fibers taken 5, 7 and 10 days after embedded in fibrin hydrogels. **C** Quantification of the number of cells migrating from the fiber to the hydrogel, considering only the ones in close proximity to the fiber length that are spreading outside, as the representative microscopic image in **D** shows (n = 12 replica fibers from 3 biologically independent experiments). **E** Cell invasion area at the final day of the assay (n = 4 replica fibers, 2 biological independent experiments). Data is presented as mean ± s.d., with *p<0.05, ***p<0.001 and ****p<0.0001 using one-way or two-way ANOVA with multiple comparisons.

After embedding, live/dead assay revealed a tendency of compartmentalized cells to migrate towards the fibrin hydrogel in both conditions, with more cells present outside near the fiber in the PEG 8% condition along the 10 days culture period (Figure 4B). Cells appear to mostly migrate the fiber’s membrane as single and occasional units, therefore we use optical microscopy to keep track on the number of cells that were in close proximity to the fiber membrane and spreading to the surrounding fibrin hydrogel as an indicator of their migration out of the fiber (Figure 4C, D). For that, we considered only the cells along the fiber length, excluding those exiting from the ends, as these regions are more prone to experimental artifacts due to the cutting process used to produce straight fibers with identical lengths. After 1 and 3 days, very few cells were observed outside the fiber-shaped compartments in both conditions. However, by day 5, a clear difference between conditions became evident, with a significantly higher number of cells in the outer region near the PEG 8% membranes – a trend that persisted throughout the 10-day culture period. Moreover, the invasion area occupied by cells in the hydrogel at the end of the assay is significantly higher in the PEG 8% condition compared to the control (Figure 4E). Together, the results indicate the ability of the explored method of manipulating interfacial membrane’s porosity in order to facilitate the migration of compartmentalized cells into surrounding tissue-like materials.

## 3. Discussion

PE complexation is a multistep process that involves an initial nucleation phase where charged groups of the polymer chains interact electrostatically displacing counter ions and water molecules, leading to further growth and organization into insoluble solid-like coacervates as water is expelled into the surrounding solution and the complex becomes denser and stable.[51] The process of PE complexation and its resultant material and structure are dependent on several factors relating to the PEs itself such as their charge density, molecular weight (MW), their intermolecular interaction, and also the solvent environment.[50,52] Recently, Liu and Tong observed the formation of interfacial complexes with a porous structure from the interaction between high MW hyaluronic acid and synthetic poly-diallyldimethylammonium (PDADMAC), while the complexation with lysine-based peptide amphiphiles led to membranes with a well-condensed non-porous structure.[20] On the other hand, Tang *et al.* reported the formation of porous membranes which are formed by spontaneous water-on-water spreading and simultaneous complexation of oppositely charged synthetic PEs.[53] The formation of such porous structures is consistent with membranes produced via non-solvent-induced or aqueous PE phase separation methods, which occur through solvent exchange and polymer precipitation, or pH-dependent charge transition behavior, respectively. [39,53,54] However, with pore sizes ranging from nanometers to just a few micrometers, these membranes are particularly well-suited for filtration applications. In here, the porous structure of membranes is affected, whether they are produced with or without the prototypical ATPS phases. Liquid-core fiber membranes produced by the complexation of EPL and ALG in an ATPS-free environment, present an increased fiber opacity suggesting a faster achievement of compact structures, and consequently show a more condensed (non-porous) membrane structure. On the other hand, porous interfacial PE membranes are created in the presence of the aqueous PEG and DEX phases. These immiscible polymer phases allow for the regulation of PE diffusion and their complexation at the interface, enabling the formation of membranes with diverse morphologies. We explore the ability of tailoring the physical properties of the resultant membranous material, by adjusting the composition of the phases involved in the PE interfacial complexation. It is well established that molecule diffusion is facilitated in low viscous environments, and that reduced ATPS polymer concentration produces solutions with lower viscosity.[25] The interaction between the PEs occurs more quickly upon the reduction of the concentration of PEG in the bath phase, due to facilitated diffusion of the EPL molecules towards the interface of the system. In fact, fibers produced in lower PEG concentration present enhanced mechanical stability, exhibiting lower swelling capacity with minimal dimensional change, and higher resistance to fiber deformation under the contractile forces of encapsulated hASCs. However, this phenomenon is not solely governed by the enhanced diffusion profile, as the estimated dry mass of the fiber membrane remains consistent across both higher and lower PEG concentrations. Instead, the membrane architecture is more openly porous in the lower viscous bath environment. This suggests that distinct molecular re-arrangements and subsequent complex network organization are playing a significant role in the observed phenomenon. The porosity of the interfacial membranes produced in PEG 8% increased 4-fold compared with the previously reported control condition using PEG at 17%. We hypothesize that upon expelling water during complexation, the ALG/EPL complexes may contract into the form of a porous network structure, which likely results in the enhanced properties observed. In PEG 17%, PE complexation remains effective, but during the dehydration step, the higher viscosity likely prevents the re-arrangement of the complexes into such an open network structure.

Additionally, PEG and DEX are both non-ionic, highly hydrophilic polymers that can act as macromolecular crowding agents.[55,56] Their strong affinity for water molecules allows them to retain hydration, while their presence in aqueous solution leads to an excluded volume effect. This effect reduces the effective volume available for other molecules, influencing their diffusion, conformation and possible interactions.[56,57] Studies using PEG as a crowding agent in complex coacervation, have shown to increase the stability of the coacervate. The role of PEG is mostly associated with a change in the conformation of the PE constituents[58] as well as the increase in the entropic dehydration driving force for the process, by extracting interstitial water from the coacervate phase without interfering with the surface hydration of the PEs.[57] In this system, decreasing the PEG concentration increases overall water availability and reduces crowding, providing more space and liberty for structural re-arrangements of the complex to occur at the interface. Another notable aspect to consider is the importance of having a more viscous inner phase and high-density contact barrier, provided by the presence of high MW DEX, to enable the generation of structurally organized porous membranes, and more stable and easily handled fiber compartments. Therefore, in general, the additional interfacial dynamics provided by the presence of both immiscible polymer phases during PE complexation at the liquid-liquid interface, was crucial to control the diffusion mechanisms, water structuring and PE interaction, necessary to manipulate the final porosity of the obtained membranes. As a consequence of the increased membrane porosity, cell-laden fibers produced in PEG 8% baths demonstrated higher potential to facilitate the migration of mesenchymal stem cells, adhered to the luminal part of the membrane, to surrounding fibrin matrices and promote its invasion.

## 4. Conclusion

In summary, we investigate how tuning the composition of immiscible concentrated polymer phases influences PE complexation at all-aqueous liquid interfaces, allowing tailoring the structural properties of the resulting membranes. Independently of the geometry of the processed interfacial material, the method remains effective and can induce different porosity degrees depending on the size of the constructs. Unlike other techniques to induce porosity in soft materials that are not amenable to be processed with encapsulated cells, the possibility to induce membrane porosity during fiber formation by simply adjusting the system composition, facilitates cell seeding of the fiber lumens in a single-step manner. The facilitated migration of mesenchymal stem cells from the highly porous fiber compartments and improved invasion of surrounding tissue-like matrices demonstrate great potential of the produced materials for tissue engineering and regeneration applications.

## 5. Methods

### Fabrication of tubular fibers

Liquid-core fibers were prepared as previously described.[35] Briefly, solutions of poly(ethylene glycol) (PEG, average MW 8000 Da, Sigma-Aldrich) and dextran (from Leuconostoc spp., MW 450,000 – 650,000 Da, Sigma-Aldrich) of varying concentration were prepared in phosphate buffer saline (PBS, Sigma-Aldrich). Then, the polyelectrolytes (PEs) sodium alginate (from brown algae, MW 120,000-190,000 g/moL, Sigma-Aldrich, 71238) and ɛ-Poly-L-lysine (EPL, derived from fermentation of Streptomyces albulus PD-1, MW ∼ 4700 g/mol, Epolyly®, Handary S.A., 0201) were dissolved separately in the dextran and PEG solutions, respectively. 2 wt% alginate and 0.75 wt% EPL was used. The pH of the PEG+EPL solution was adjusted to ∼7.2, after the complete dissolution of EPL. The fibers were formed by dispensing a solution of dextran and alginate using a syringe pump (Harvard Apparatus) with a flow rate of 0.2 mL/min, in a larger volume bath phase of PEG and EPL. The petri dish containing the bath solution is displaced during extrusion of the dispensing phase, to allow the formation of threads where interfacial PE complexation occurs. The produced structures are segmented using a spatula, to enable the generation of straight fibers with control over their length. After 2 minutes, the complexation is interrupted by removing the bath phase and performing 3 washing cycles with PBS. A 2-minute complexation time was used for the tested conditions, unless otherwise mentioned. To assess the opacity of the membranes, the mean gray value was determined from images taken in an optical microscope (Zeiss PrimoVert), using ImageJ.

### Fabrication of spherical capsules

Spherical capsules were produced as described previously.[28] Briefly, millimetric-sized capsules were fabricated by manual dropwise addition of the dispensing phase containing dextran 15 wt% and ALG 2 wt%, using a 25G needle, to a 30 mL bath of PEG 17 wt% and EPL 0.75 wt%, stirring at 300 rpm. Using the same phase compositions, the micro-sized capsules were produced through electrohydrodynamic atomization using a EF100 Desktop Needle System Electrospinning/Electrospraying equipment. The dispensing phase was sprayed in the collecting bath phase under stirring at 300 rpm, using a 22 G needle, a flow rate of 10 mL/h and 10 kV of voltage. In both cases, after complexation occurred for 2 minutes, the structures were washed with PBS three times.

### Swelling ratio

The expansion of the fibers while immersed in PBS was monitored overtime. Tubular fibers made in bath solutions with different PEG concentrations were produced as previously described using the syringe pump, and images were acquired using a microscope (Zeiss Primovert) to determine their external diameter using ImageJ. The swelling ratio (ɛ) was calculated by the change in size as follows:

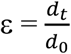

where d_0_ represents the original diameter of the fibers in the bath solution, and d_t_ the diameter of the fibers against PBS at different timepoints.

### Estimated dry weight

Lyophilization was used to estimate the dry weight of the produced fiber-shaped soft compartments. For that, 0.5 cm long fibers from each condition were produced using the pump, were left to equilibrate in the PBS washing solution, and further thoroughly washed with distilled water to remove potential effects of salt in the final weight. Also, they were ruptured to remove the inner liquid. A total number of 12-20 fibers were placed together in an Eppendorf tube®, frozen overnight at −80°C, and then freeze-dried. The fiber dried mass was estimated from the total weight after the lyophilization, using a total of 8-9 tube replicas for each condition, from 3 independent experiments.

### Membrane porosity characterization

To investigate the porous features of the interfacial membranes, confocal fluorescence microscopy was used. For that, washed fibers were incubated overnight at 4°C in a solution of fluorescein 5(6)-isothiocyanate (FITC, Sigma-Aldrich) in PBS (1:50, from a 1 mg/mL FITC solution prepared in DMSO). After washing with large amounts of PBS to remove unreacted FITC, z-stacks were acquired in stained fibers under a confocal microscope (Zeiss LSM900, Carl Zeiss Microscopy, Germany), and images were analyzed using ZEN 3.0 software. The porosity and pore size was determined by the % of total pore area and individual areas of the most representative pores, respectively, from 2D maximum projection images, using ImageJ software.

### Scanning electron microscopy

SEM analysis was performed to analyze the morphology of the inner and outer surfaces of the membranes with different porosity. Liquid-core fibers were dehydrated with ethanol solutions following a concentration gradient of: 30% (v/v), 50% (v/v), 70% (v/v), 80% (v/v), 90% (v/v), 96% (v/v) and 100% (v/v), by immersing them in these solutions for 15 min. To visualize the inner surface of the membrane, cross-sections were made in the fibers using a scalpel blade, prior to dehydration. Samples were then imaged using a Hitachi SU3800 Scanning Electron Microscope, with an acceleration voltage of 5 kV.

### Cell culture and encapsulation

Human mesenchymal stem cells from the adipose tissue (hASCs) were purchased from Lonza (PT-5006). hASCs were cultured in α-minimum essential medium (α-MEM; Gibco), supplemented with 2.2 g/L sodium bicarbonate (Sigma-Aldrich), 10% (v/v) fetal bovine serum (FBS, Gibco) and 1% (v/v) antibiotic/antimycotic solution (ATB, Gibco - 10,000 U/mL of penicillin, 10,000 μg/mL of streptomycin, and 25 μg/mL of Amphotericin B) at pH 7.4. Cells were used in passages between 6 and 8. C166-GFP mouse endothelial cell line (from the yolk sac) was kindly given by Dr. Enrique Martínez-Campos from ICTP-CSIC (Madrid, Spain). These cells were cultured with Dulbecco’s Modified Eagle Medium high glucose (ThermoFisher Scientific), 3.7 g/L sodium bicarbonate, 10% FBS, 1% ATB. For the selection of GFP-expressing cells, G418 antibiotic was used (0.2 mg/mL). Cells were used in passage 4.

For cell encapsulation procedures, both aqueous phases were sterilized by filtration and UV irradiation, as previously described.[28,35] Sodium alginate NOVATACHTM MVG GRGDSP (GRGDSP-coupled high MW alginate, NovaMatrix, Norway) was added to the dispensing phase at a concentration of 0.5 wt% to promote cell adhesion. Cell suspensions were obtained after detaching the cells with trypsin-EDTA solution (ThermoFisher Scientific), and further centrifugated and mixed with the dispensing phase at a density of 10 million cells/mL. A syringe-needle manual injecting method was used to prepare the cell-laden fibers to facilitate the process in a sterile environment. Bath phases of PEG at concentrations of 17% and 8% with EPL 0.75%, were used to prepare the fiber-shaped membranes with different degrees of porosity. After 2 min of complexation, the bath phase was removed and DPBS was added (×3). Fibers of identical length were then transferred to a DPBS solution (30 mL) and finally to the appropriate cell culture medium in well plates. Cells and cell-laden compartments were maintained in incubators with controlled temperature (37 °C) and 5% CO_2_.

### Cell-mediated fiber deformation

The deformation of the fibers was monitored throughout the culture period by capturing images at specific timepoints using an inverted microscope (Zeiss PrimoVert) with a coupled camera. Fiber area contraction and bending angles were determined using ImageJ. The bending angle is defined by θ, which is the measure of the vertex angle made from three points: one corresponding to the center of the fiber at the bending region, and the other two corresponding to equidistant middle points of farer regions of the fiber, in all samples. To inhibit actomyosin contractility, (−)-blebbistatin (Sellek Chemicals, S7099) was supplemented in the cell culture medium at a final concentration of 25 μM.

### Cell viability assays

The cell viability of cell-laden fibers was analyzed by live/dead assay (Invitrogen, USA) and Alamar Blue® Cell Viability assay (Thermo Fisher Scientific). Both assays were performed in accordance with manufacturer instructions. To analyze live and dead cells at pre-determined timepoints, fibers were incubated in cell culture medium with propidium iodide (PI) (1:1000, Thermo Fisher Scientific) and Calcein-AM solution (1:500, Thermo Fisher Scientific) during 10 min at 37°C. Fibers were washed with culture medium and examined in an upright widefield fluorescence microscope (Axio Imager M2, Carl Zeiss, Germany). The AlamarBlue® assay was used to access metabolic activity of encapsulated cells. Cell-laden fibers with approximately 0.5 cm long were placed in 48 well-plates, with one fiber per well. AlamarBlue^TM^ reagent was added to the cell culture medium (1:10), and incubated for 8 hours. Fluorescent measurements (λ_excitation_: 540 nm, λ_emission_: 600 nm) were performed in a Synergy HTX microplate reader using a 96-well black-clear bottom plate. A representative number of five fibers per independent experience were used, and fibers without cells were used as controls.

### Cell migration assay in fibrin matrices

To evaluate the cell migration capacity from liquid-core tubular compartments with different membrane porosity to surrounding matrices, cell-laden fibers were embedded within fibrin hydrogels. A 4-day period of pre-incubation was used, to promote the adhesion and organization of the cells within the produced materials. After that, fibers were placed in cell culture chamber slides with square-shaped wells (µ-Slide 8 Well high, ibidi, 80806), and a solution of fibrinogen (from human plasma, Sigma-Aldrich, F3879) at 4 mg/mL in cell culture medium with aprotinin (from bovine lung, Sigma-Aldrich, A4529) at 25 µg/mL was added, followed by a thrombin (from human plasma, Sigma-Aldrich, T6884) solution at 2 U/mL in PBS (mixing ratio of 1:1). The slides were incubated at 37°C for 30 minutes for fibrin gelation and media was added on top. To provide a humidified environment and prevent rapid evaporation, the slides were maintained into petri dishes with a small amount of PBS. Media was changed every 1-2 days.

### Statistics and data presentation

All statistical analysis was performed using GraphPad Prism 8.0.2. Statistical significance was assessed by one or two-way analysis of variance (ANOVA) with Tuckey’s or Sidak’s multiple comparison tests, where a p value < 0.05 was considered significant. A line was denoted between significant groups, straight line comprising all groups bellow the line, and unlabeled groups in these graphs have no statistical significance. All data are presented as mean ± standard deviation, except for the violin plots that represent the pore size distribution using density curves. N replicate values and biological independent experiments are described in figure captions. Schematics were designed with Adobe Illustrator.

## Acknowledgements

This work was developed within the scope of the project CICECO-Aveiro Institute of Materials, UIDB/50011/2020, UIDP/50011/2020 & LA/P/0006/2020, financed by national funds through the FCT/MCTES (PIDDAC). It was financially supported by the European Research Council grant agreement ERC-2019-ADG-883370 (project REBORN). Also, the Programa Operacional Competitividade e Internacionalização, in the component FEDER, supported this research by national funds (OE) through FCT/MCTES, in the scope of the project ‘CellFi’, PTDC/BTM-ORG/3215/2020. Raquel C. Gonçalves acknowledges financial support received from FCT trough individual grant 10.54499/2021.07435.BD. We sincerely thank Ana Santos-Coquillat for kindly providing the GFP-C166 cells used in this study.

## Data availability Statement

The data that support the findings of this study are available from the corresponding author upon request.

## Supporting Information

**Figure S1.**
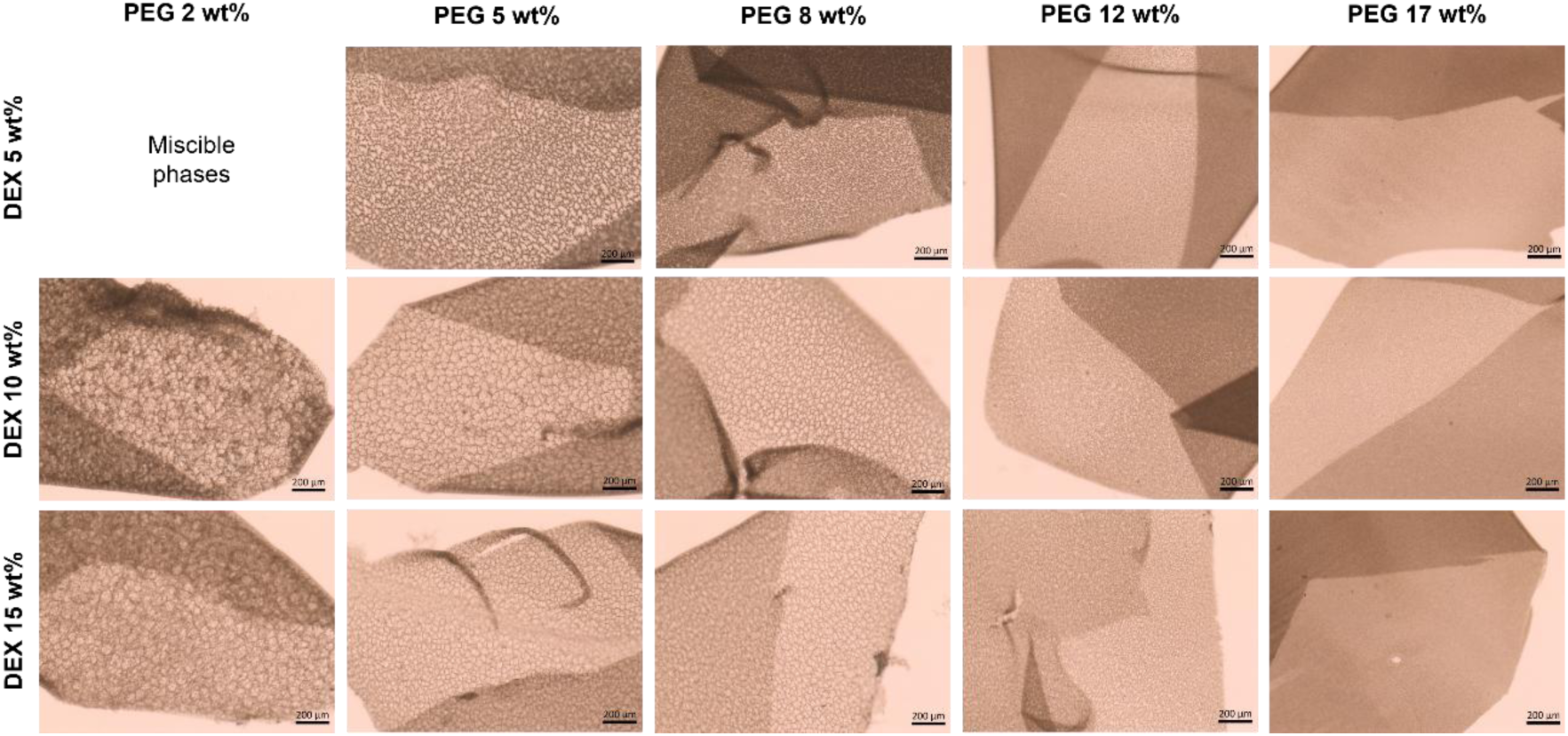
Effect of polymer concentration on membrane morphology. Representative brightfield microscope images of fiber membranes produced using prototypical polymer systems with variable PEG and DEX concentration, showing that decreasing the PEG concentration induced the formation of a pore-like morphology at the interfacial membrane, while the variation of DEX concentration didn’t induce highly visible effects. The structures were washed with PBS after a 2 minutes complexation time, and were ruptured to allow the visualization of the membrane morphology. Fibers formed with the lowest polymer concentrations were not considered since those concentrations did not allow phase separation, therefore the phases were not immiscible.

**Figure S2.**
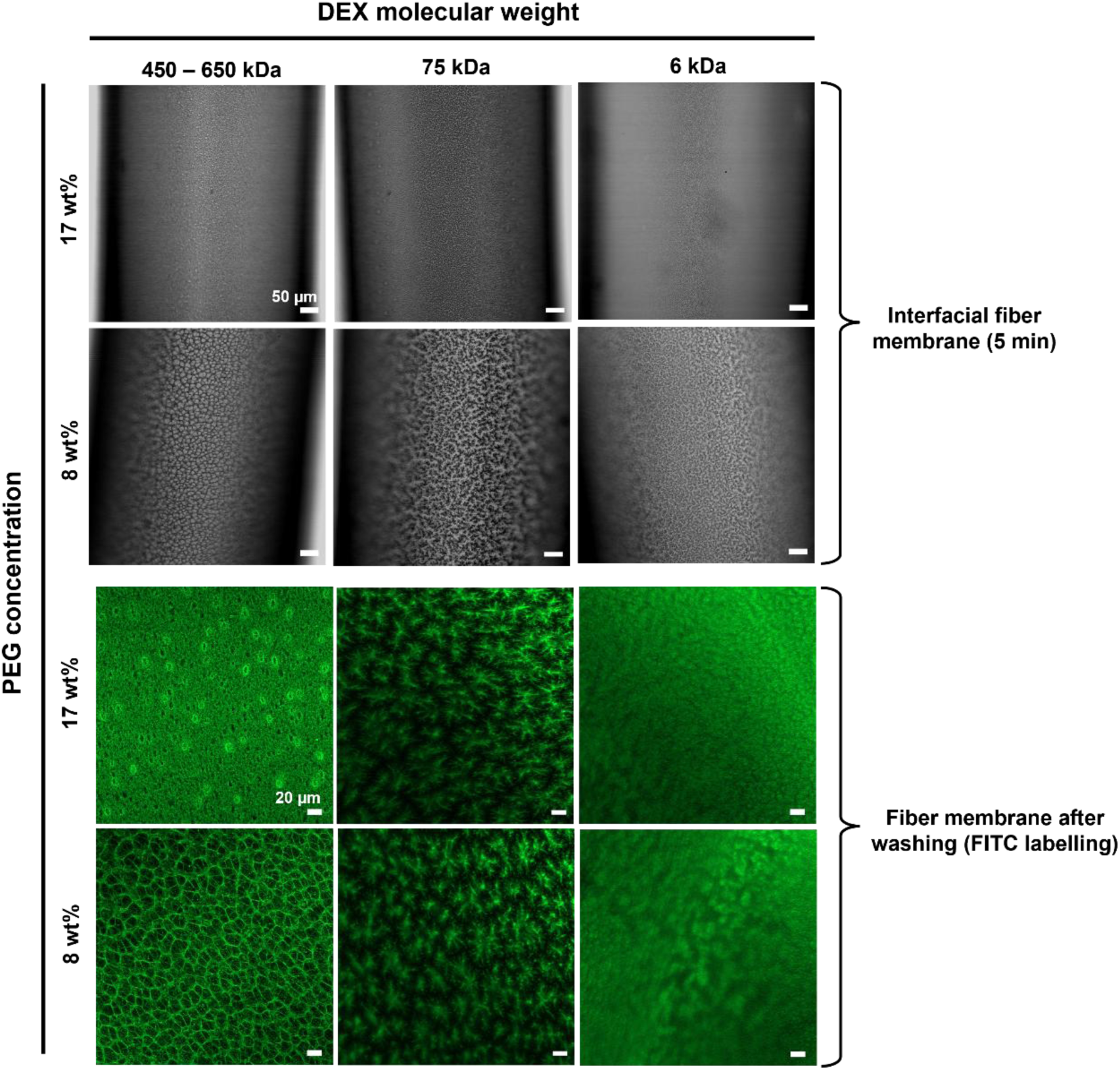
Effect of DEX molecular weight on membrane morphology. On the top, representative images of interfacial fiber membrane within 5 minutes of contact between the dispensing phase containing different DEX with different molecular weight (450-650 kDa, 75 kDa and 6 kDa) and the bath phase containing PEG 8 kDa with distinct concentration – 17 wt% and 8 wt%. On the bottom, are representative confocal images of FITC-labelled fiber membranes, washed with PBS after 2 minutes complexation time. The images show that higher DEX molecular weights allow the formation of more interesting membrane morphologies, and that decreasing PEG concentration continues to promote a higher interspaced complex network.

**Figure S3.**
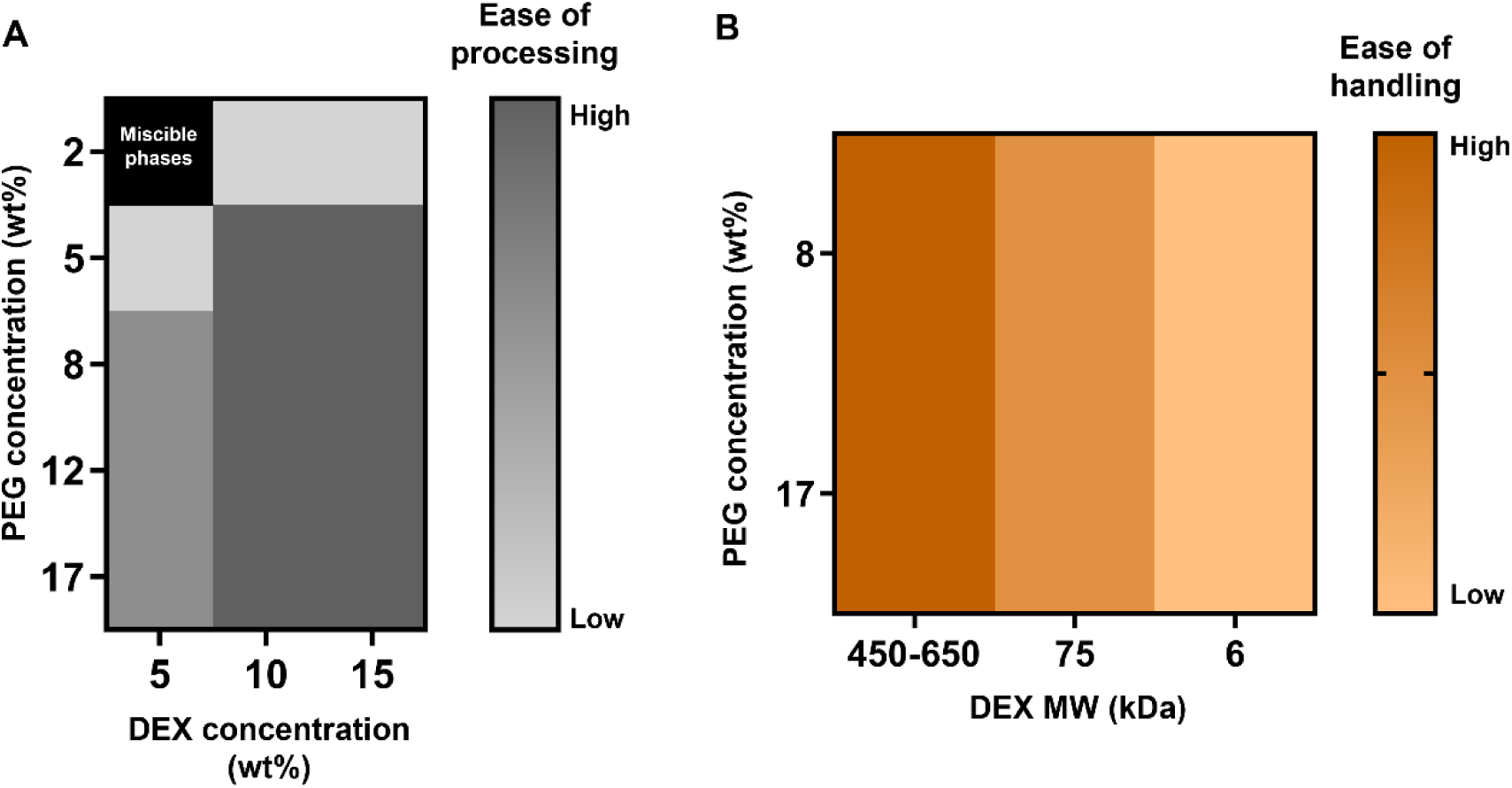
Qualitative evaluation of fiber processing and handling properties with different prototypical ATPS composition. **A** Heatmap representing the level of ease of processing of fibers produced with different polymer concentrations. Lower polymer concentration – DEX 5 wt% and PEG 2 wt% - mainly difficulted the processing of the fibers due to either low dispensing phase density or low bath viscosity, respectively. **B** Heatmap of the handling capacity of fibers made with dextran with different molecular weight (MW). As the molecular is reduced, it is more difficult to handle the fibers mainly because they tend to stick together, increasing their risk of rupture.

**Figure S4.**
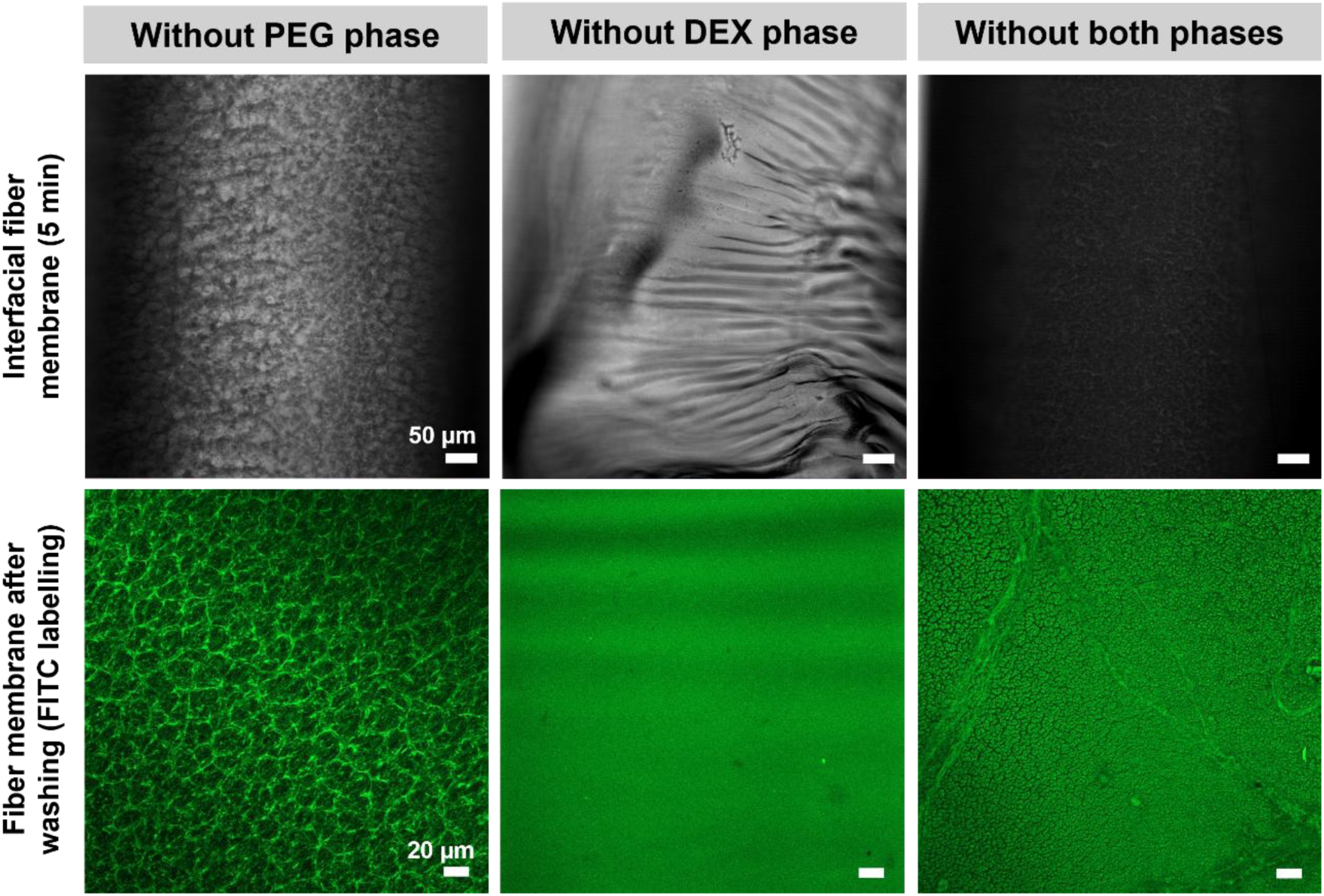
Control conditions to evaluate the effect of the distinct polymer phases on the interfacial membrane morphology. Interfacial membrane formed without the presence of the PEG phase, DEX phase and both the immiscible phases. Images on the bottom are of the washed and FITC-labelled fiber structures. While removing only the PEG from the fiber formulation, produced membranes with similar pore-like morphology although with a more closed network, the removal of the DEX phase produced interfacial membranes with a more homogeneous structure. The removal of both phases produced highly dense membranes with no morphological organization.

**Figure S5.**
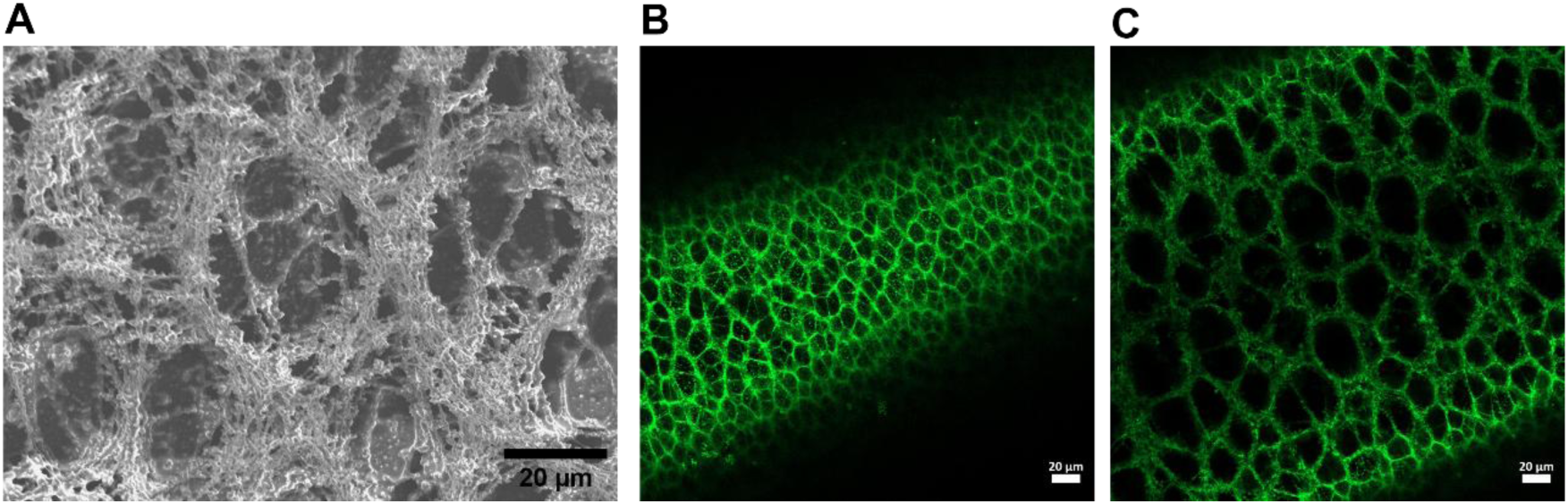
Characterization of interfacial membranes produced in PEG 8% baths. **A** Scanning electron micrograph of a PEG 8% membrane in a closer view, showing the presence of a highly filamentous microstructure. Representative confocal florescence images of the porous nature of an outer in **B** and inner in **C** layers of the membrane, respectively, indicating a tendency to generate larger pores at the luminal part of the liquid-core fiber.

**Figure S6.**
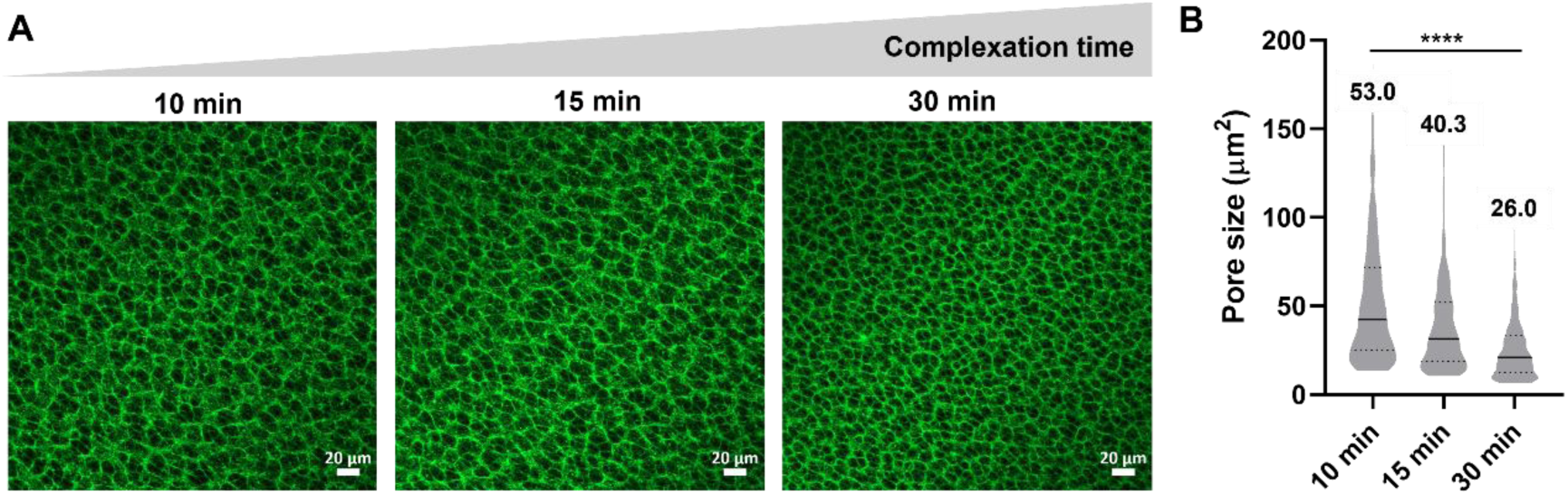
Versatility of membrane porosity by exploring complexation time. **A** Representative images of FITC-labelled membranes produced with varying complexation times – 10, 15 and 30 minutes – using a bath containing PEG at 8%. **B** Quantification of pore size of membranes with the different complexation times, showing a decrease in pore size as the complexation time increases.

**Figure S7.**
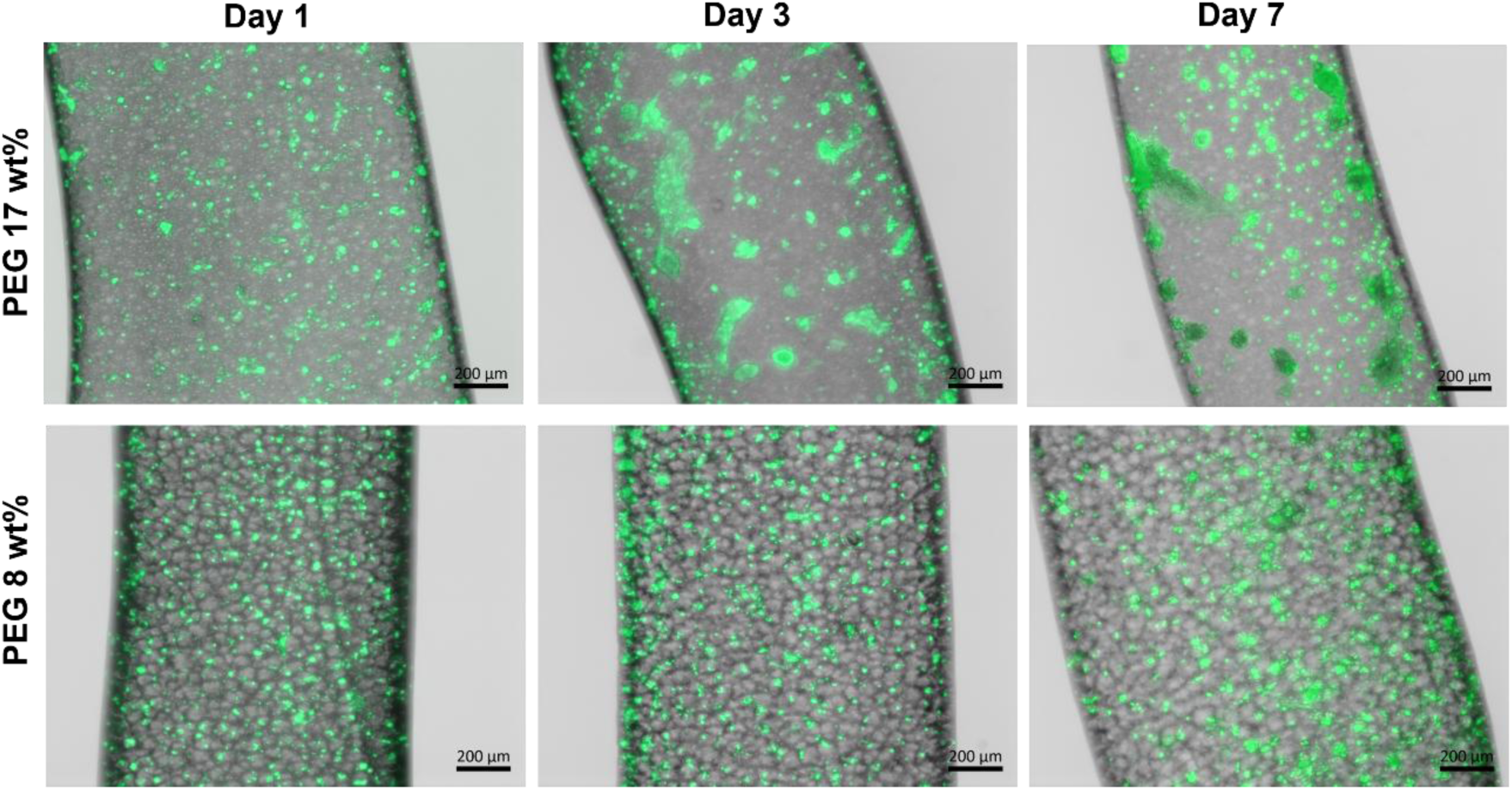
Organization of mouse endothelial cells. Representative fluorescence micrographs of GFP-expressing C166 mouse endothelial cells encapsulated within fiber compartments, showing a higher tendency to undergo cell aggregation in the PEG 17% control condition, compared to the more uniformly distributed cells in the PEG 8% condition.

**Figure S8.**
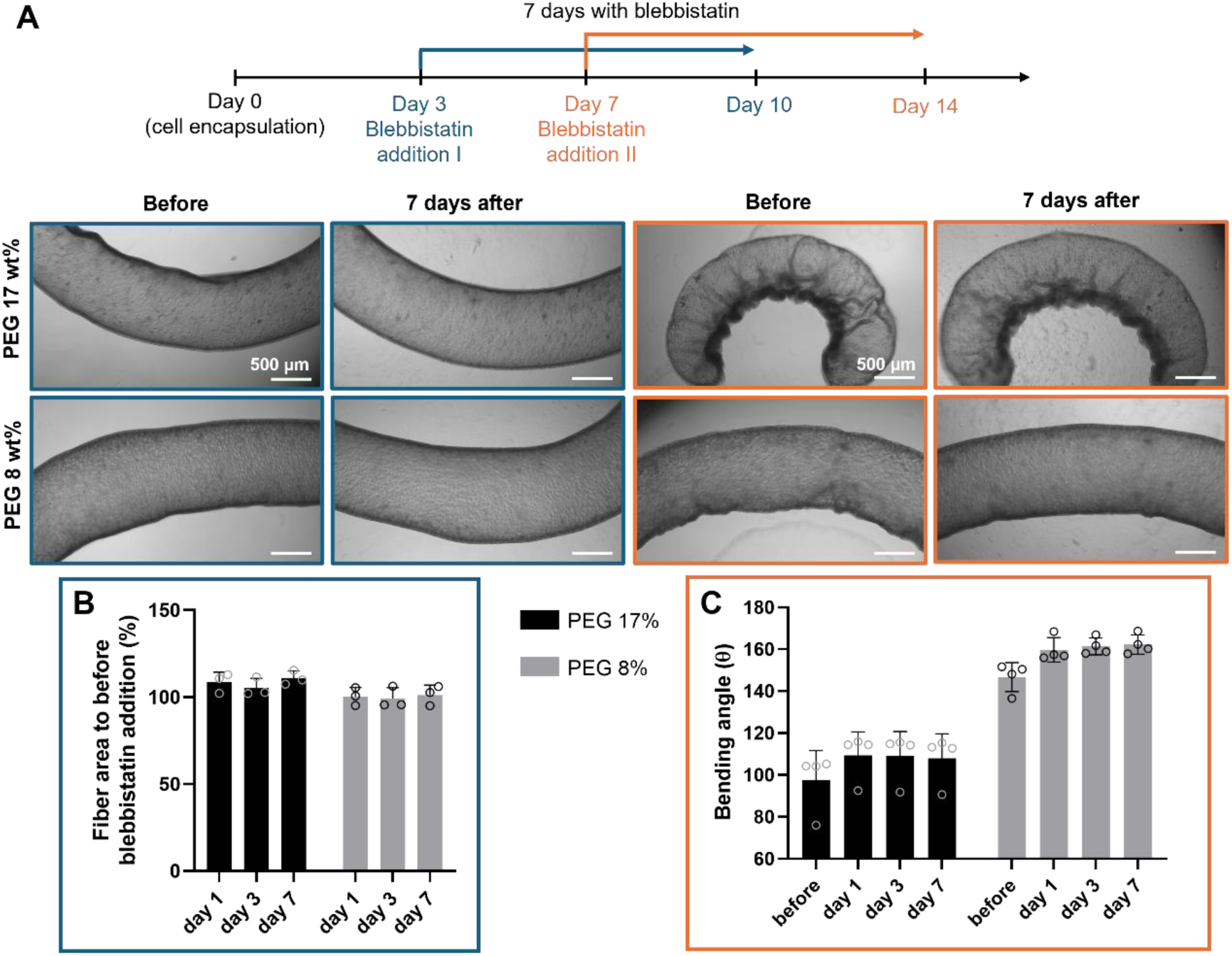
Fiber deformation in the presence of blebbistatin. **A** Timeline of the strategy used to study the effects of blebbistatin addition as a prevention of fiber contraction (at day 3), or as a possible way to revert already deformed fibers (at day 7). On the bottom there are representative optical microscope images of the fibers before the addition of blebbistain, and 7 days after. **B** Quantification of fiber area as percentage of the original area before blebbistatin addition, demonstrating negligible variation of fiber area over the 7 days of culture in both conditions, which indicates that blebbistatin prevented fiber contraction (n = 3 replica fibers, 1 biologically independent experiment). **C** Quantification of the bending angle of 7-days deformed fibers before and after the treatment with blebbistatin, showing a slight increase of the angle after 1 day, although not statistically significant (n = 4 replica fibers, 1 biologically independent experiment). Data is presented as mean ± s.d.

**Video S1.** Z-stack throughout the thickness of liquid-core fiber membranes produced in PEG 8% baths, after washing and labelling with FITC.

